# Creator 1.0: Phylogeny-Guided Stagewise Pretraining for Plant Genome Foundation Models across 1,000 Species

**DOI:** 10.64898/2026.10.22.761197

**Authors:** Lin Yang, Sirong Jiang, Yubo Li, Meiling Zou, Yu Xia, Lu Wang, Yingdong Wu, Hao Wang, Wei Xie, Zhiqiang Xia

**Affiliations:** School of Breeding and Multiplication (Sanya Institute of Breeding and Multiplication)/ Hainan Seed Industry Laboratory/ College of Tropical Agriculture and Forestry/ State Key Laboratory of Tropical Crop Breeding, Hainan University, Sanya, Hainan, 572025, China; Sanya Hyper Gene Biotechnology, Sanya, Hainan, 572025, China

**Keywords:** plant genomic language model, phylogeny-guided stagewise pretraining, checkpoints, cross-species evaluation

## Abstract

Genomic language models have advanced DNA sequence analysis, but their applicability to plant-specific tasks and representation changes during pretraining remain poorly characterized. Here, we present Creator 1.0, a 2-billion-parameter parameter plant genome foundation model pretrained on 1,000 plant genomes using phylogeny-guided stagewise pretraining (PGSP). Training comprised 83 consecutive stages and produced 52 cumulative checkpoints associated with distinct plant orders. On GENEB, the stage 83 checkpoint ranked first among 46 models across 10 plant-related tasks, with a mean Matthews correlation coefficient (MCC) of 0.565, and fourth across all 100 tasks. Evaluation of 52 checkpoints on six plant lncRNA tasks identified different optimal stages under full-data, 10-shot and 1-shot settings, corresponding to stages 71, 72 and 29, respectively. Checkpoint performance varied across tasks, without consistent advantages for corresponding plant orders. These findings demonstrate the competitive performance of Creator 1.0 and highlight the potential of PGSP to provide task-adaptable checkpoints for plant genomic applications.

## 1. Introduction

Genomic language models learn sequence representations through self-supervised pretraining on large-scale DNA sequence datasets and have been applied to genomic function prediction, regulatory element identification and genetic variant effect prediction ^1-4^. Studies such as Nucleotide Transformer and AgroNT have demonstrated that pretrained models can provide transferable sequence representations across species and downstream tasks^5-7^. As pretraining datasets expand to encompass broader phylogenetic diversity ^8^, an important challenge in developing and evaluating multispecies genomic models is to capture sequence features shared across species while preserving genomic characteristics specific to different plant lineages.

The performance of genomic language models is influenced by pretraining data, evaluation tasks and downstream adaptation strategies, making results from different studies difficult to compare directly ^9^. BEND and DART-Eval provide complementary frameworks for evaluating genomic models ^10, 11^, whereas GENEB evaluates model performance using frozen representations and linear probing across 100 DNA sequence classification tasks spanning 13 functional categories, including 10 plant-related tasks ^9^. GENEB therefore provides a standardized framework for comparing plant-specific performance and generalization across tasks. However, existing evaluations of multispecies genomic models have primarily focused on final pretrained models ^12^, leaving changes in learned representations across pretraining stages and their task-specific performance differences insufficiently characterized.

Here, we developed Creator 1.0, a plant genome foundation model trained using phylogeny-guided stagewise pretraining (PGSP), in which genomic sequences from 1,000 plant species were organized according to a fixed phylogenetic tree. Model checkpoints were retained throughout successive training stages to enable systematic evaluation of stage-dependent performance. We first compared the GENEB evaluation results of six representative Creator 1.0 checkpoints with published benchmark results from 40 models, assessing their performance on plant-related tasks and across all 100 benchmark tasks. We then systematically examined performance trajectories and inter-task correlations across 52 cumulative checkpoints labeled by plant taxonomic order, using six plant long non-coding RNA (lncRNA) classification tasks under standardized evaluation conditions.

## 2. Results

### 2.1. Model architecture and stage-specific checkpoints

The pretraining corpus comprised approximately 1.11 Tbp of genomic sequences from 1,000 plant species, representing 958 genera, 199 families and 52 orders. PGSP organized the corpus into 83 consecutive training stages according to a fixed phylogenetic tree. Model parameters were carried forward between successive stages, with checkpoints and associated training metadata retained throughout pretraining (Fig. 1). Creator 1.0 is a causal Transformer comprising 30 layers and approximately 2.014 billion parameters. DNA sequences were tokenized into non-overlapping 3-mers, with a maximum input length of 32,768 tokens. Each checkpoint represents the cumulative state of the model after preceding training stages rather than an independently trained model specific to a given plant order. Of these checkpoints, 52 were associated with plant order labels.

**Fig 1.**
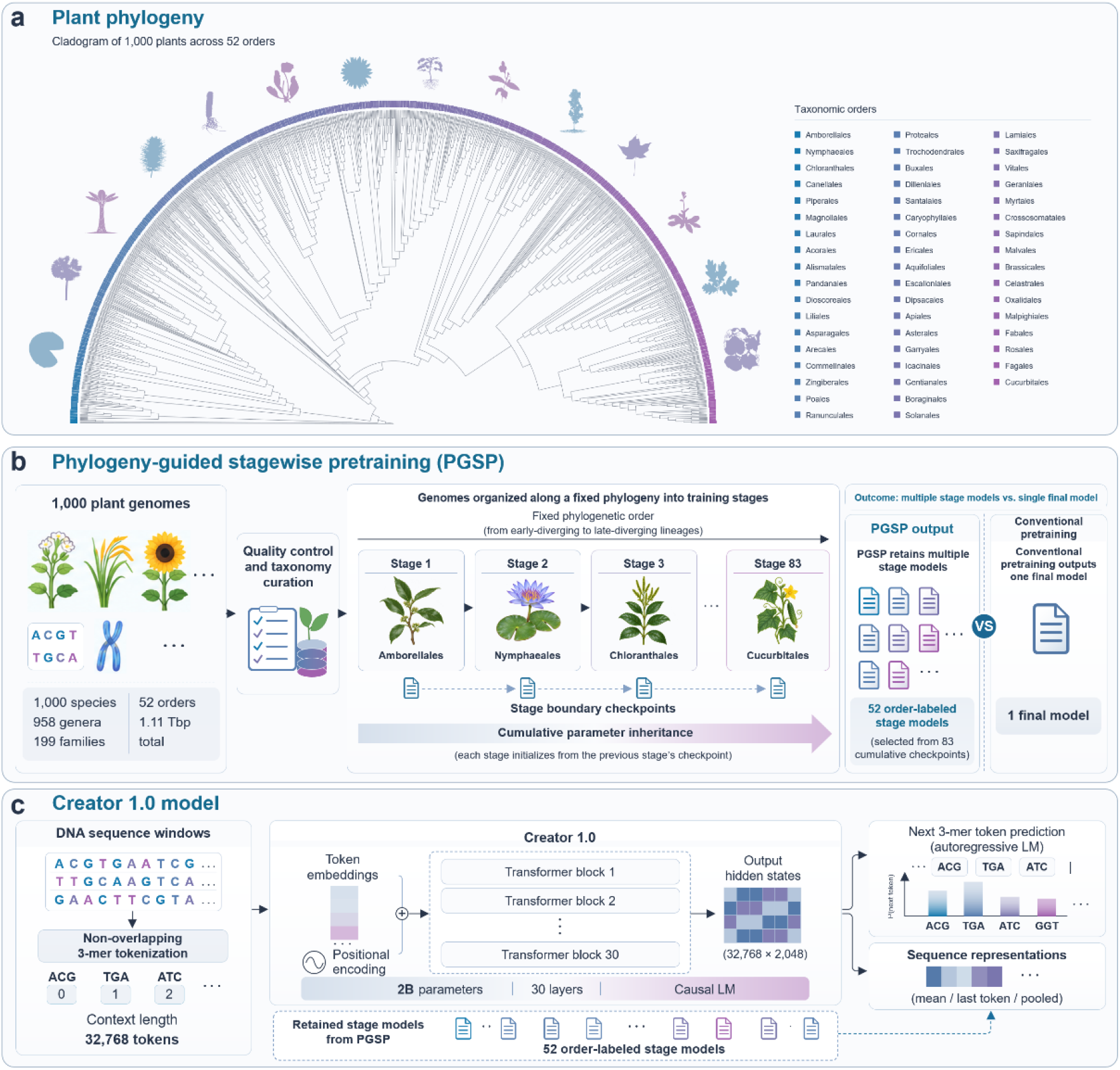
Phylogeny-guided stagewise pretraining and model architecture. a, Phylogenetic relationships among 1,000 plant species spanning 52 orders, illustrating the taxonomic diversity of the pretraining corpus. b, Overview of phylogeny-guided stagewise pretraining (PGSP). Plant genomes were organized into 83 consecutive training stages according to a fixed phylogenetic tree. Each stage inherited parameters from the preceding checkpoint, producing multiple cumulative models, including 52 checkpoints associated with distinct plant orders, in contrast to conventional pretraining that typically retains a single final model. c, Architecture of Creator 1.0. DNA sequences were encoded into non-overlapping 3-mers and processed by a 2-billion-parameter causal Transformer comprising 30 layers, a hidden dimension of 2,048 and a maximum context length of 32,768 tokens. The model supports autoregressive next-token prediction and sequence representation extraction, with multiple stage-specific checkpoints retained for downstream applications.

### 2.2. Creator 1.0 achieves the highest mean performance on GENEB plant-related tasks

GENEB includes 10 plant-related tasks covering promoter identification and DNA methylation in *Arabidopsis thaliana*, PRO-seq-related classification in cassava, and lncRNA classification across six plant species. We compared six Creator 1.0 checkpoints with 40 publicly available reference models. Under the full-data setting, the stage 83 checkpoint achieved a mean MCC of 0.565 across the 10 plant-related tasks, ranking first among all 46 models (Fig. 2). All six Creator checkpoints ranked within the top 10, with four among the top five. The mean MCC of stage 83 exceeded those of PlantCaduceus and AgroNT by 0.112 and 0.126, respectively, and its performance was higher than both reference models on all 10 tasks. Stage 83 achieved MCC values of approximately 0.90–0.92 on the *A. thaliana* promoter prediction task and 0.41–0.57 across the six lncRNA classification tasks, indicating substantial variation in predictive performance across task types.

**Fig 2.**
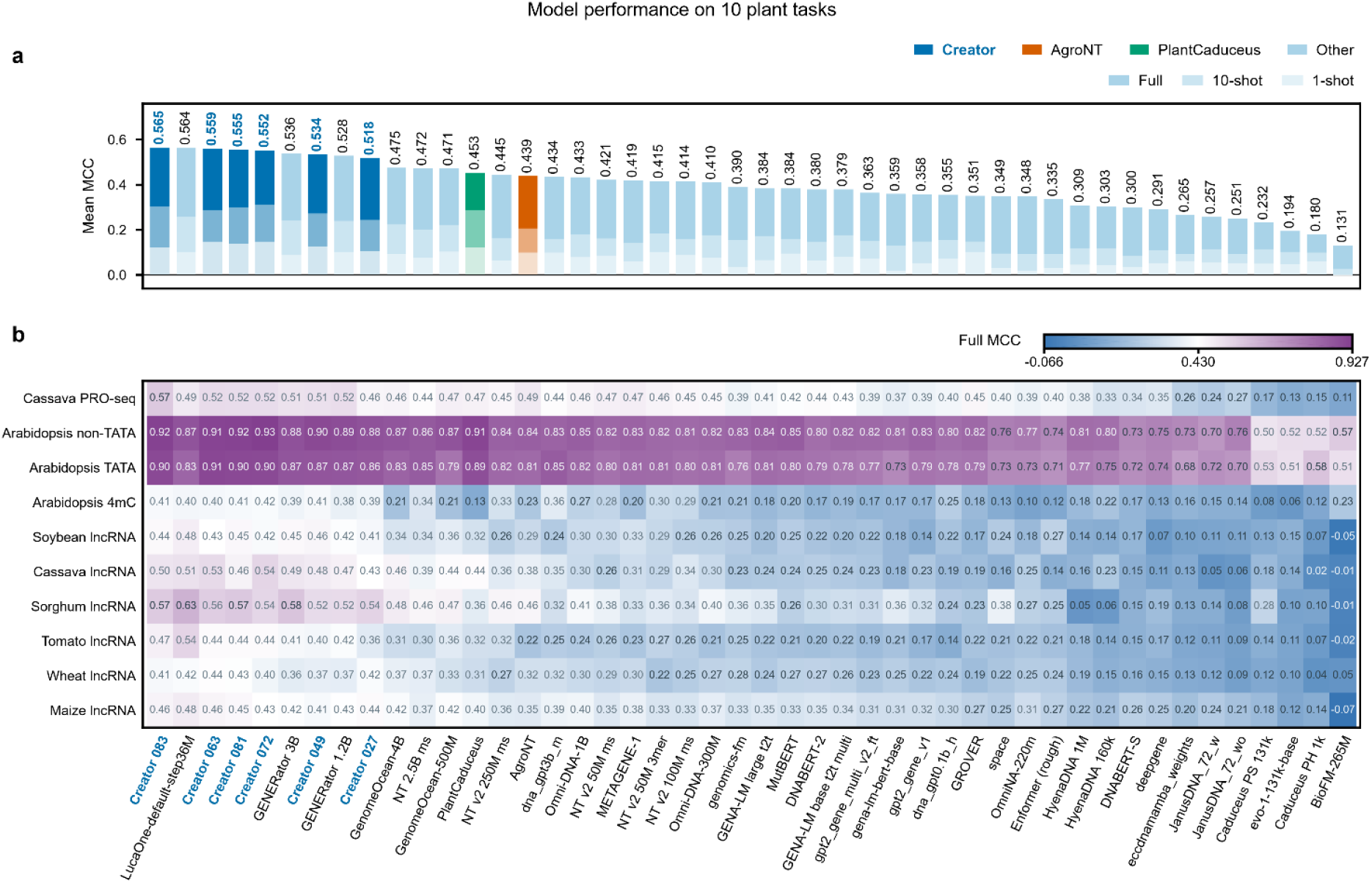
Performance of 46 models across 10 plant-related genomic tasks. a, Arithmetic mean MCC across 10 plant-related tasks for 46 models under full-data, 10-shot and 1-shot settings. Overlaid bars share a common zero baseline, with the endpoint of each bar indicating the mean MCC for the corresponding evaluation setting. b, Full-data MCC values for each model across the 10 tasks. Models in both panels are ordered by descending arithmetic mean of unrounded full-data MCC values across the 10 tasks. Creator model names are highlighted in bold blue. Individual MCC values are annotated in the heatmap, with blue, white and purple corresponding to the observed minimum, arithmetic midpoint and observed maximum of the displayed range, respectively.

### 2.3. Creator 1.0 maintains competitive representation performance across the full GENEB benchmark

Across all 100 GENEB tasks, the mean full-data MCC of the six evaluated Creator checkpoints increased from 0.5458 at stage 27 to 0.567 at stage 83, corresponding to an improvement in ranking from 13th to fourth among 46 models. The highest-scoring model, GENERator-eukaryote-3b-base, achieved a mean MCC of 0.5979 (Fig. 3). Under the 10-shot and 1-shot settings, stage 83 achieved mean MCC values of 0.3106 and 0.1439, ranking seventh and ninth, respectively. Among 86 explicitly identified non-plant tasks, stage 83 outperformed the published GENEB reference scores of AgroNT and PlantCaduceus on 77 and 85 tasks, respectively. Another checkpoint, stage 81, achieved the highest MCC among the 46 models on three 4mC classification tasks involving *Drosophila, Geobacter pickeringii* and *Geoalkalibacter subterraneus*, with scores of 0.4935, 0.2943 and 0.2183, respectively. These results indicate that Creator 1.0, despite being pretrained exclusively on plant genomic sequences, can achieve competitive performance on selected non-plant genomic tasks.

**Fig 3.**
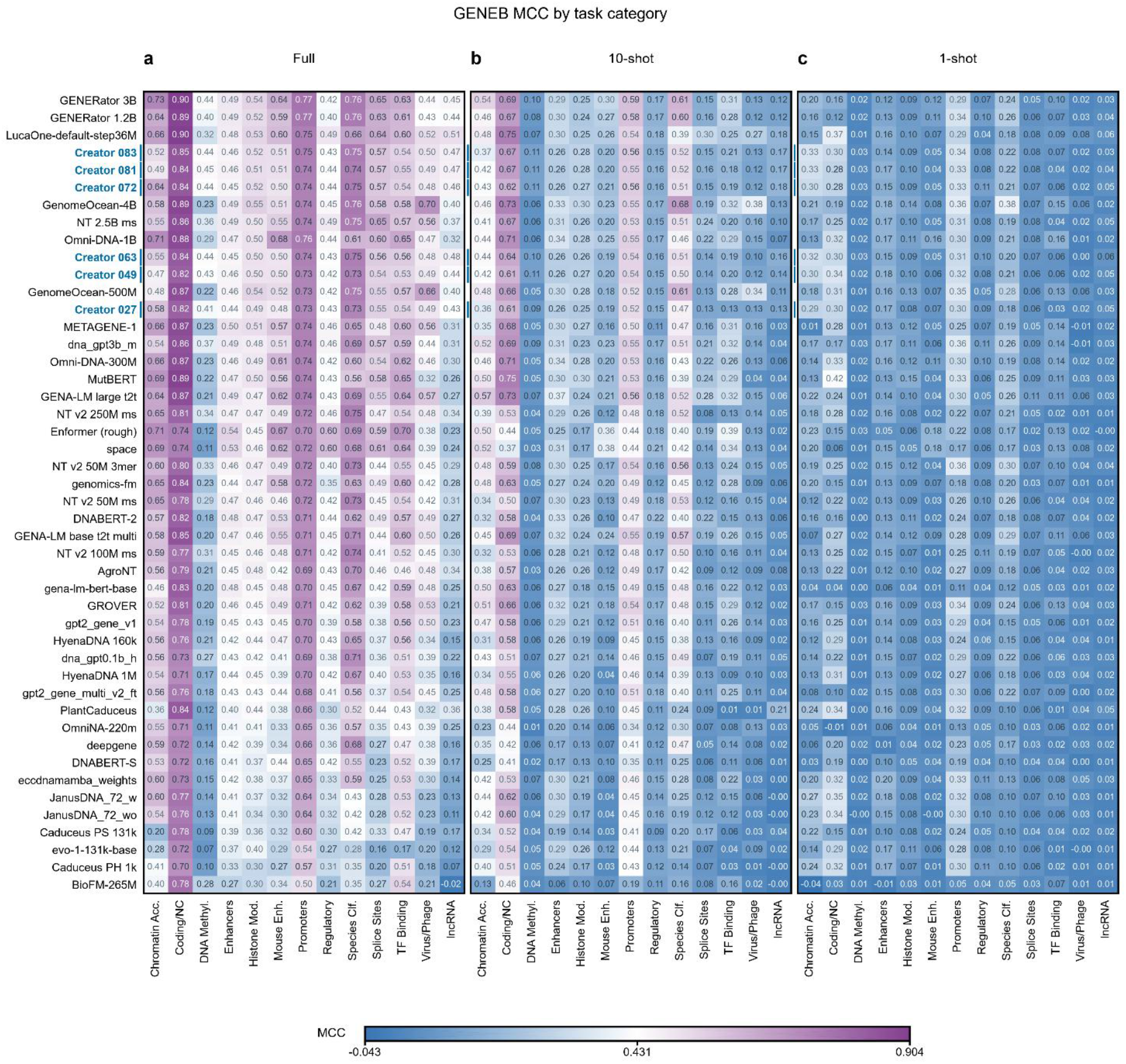
Performance of 46 models across 13 GENEB functional categories under three evaluation settings. a–c, Mean MCC values for 46 models across 13 GENEB functional categories under full-data, 10-shot and 1-shot settings, respectively. All panels use the same model order, determined by descending arithmetic mean full-data MCC across all 100 tasks. Each heatmap cell represents the arithmetic mean MCC of tasks within the corresponding functional category. The six Creator checkpoints are highlighted using blue model labels and row-side markers. All three panels share a common color scale.

Overall rankings were sensitive to task composition and score aggregation. Stage 83 ranked 17th, fifth and 21st in mean full-data MCC across human (47 tasks), yeast (20 tasks) and mouse (6 tasks) datasets, respectively. When the aggregation method was changed from equal weighting of all 100 tasks to equal weighting of the 13 functional categories, its overall ranking decreased from fourth to eighth. Although the mean full-data MCC across all 100 tasks increased with stage number among the six selected checkpoints, stage 83 achieved the highest score in only 6 of the 13 functional categories within this subset. Thus, overall benchmark rankings do not fully capture performance differences across species or functional categories.

### 2.4. Stage-dependent representation differences across plant lncRNA tasks

We evaluated 52 Creator checkpoints on six plant lncRNA classification tasks and independently reran AgroNT and PlantCaduceus as baseline models under the same local evaluation framework. Among the 52 checkpoints, the highest observed mean MCC across the six tasks was achieved at stage 71 (0.4863), stage 72 (0.1772) and stage 29 (0.0624) under full-data, 10-shot and 1-shot settings, respectively. None of these corresponded to the final training stage. Stage-dependent performance differences across individual tasks under the full-data setting are shown in Fig. 4. Stage 71 achieved a higher mean full-data MCC than locally evaluated PlantCaduceus (0.3665) and AgroNT (0.3377). Under the 10-shot setting, however, the highest-scoring Creator checkpoint (stage 72) performed below PlantCaduceus (0.2341). Under the 1-shot setting, the performance difference between stage 29 and PlantCaduceus (0.0576) was small. No single checkpoint consistently achieved the highest performance across all supervision settings.

**Fig 4.**
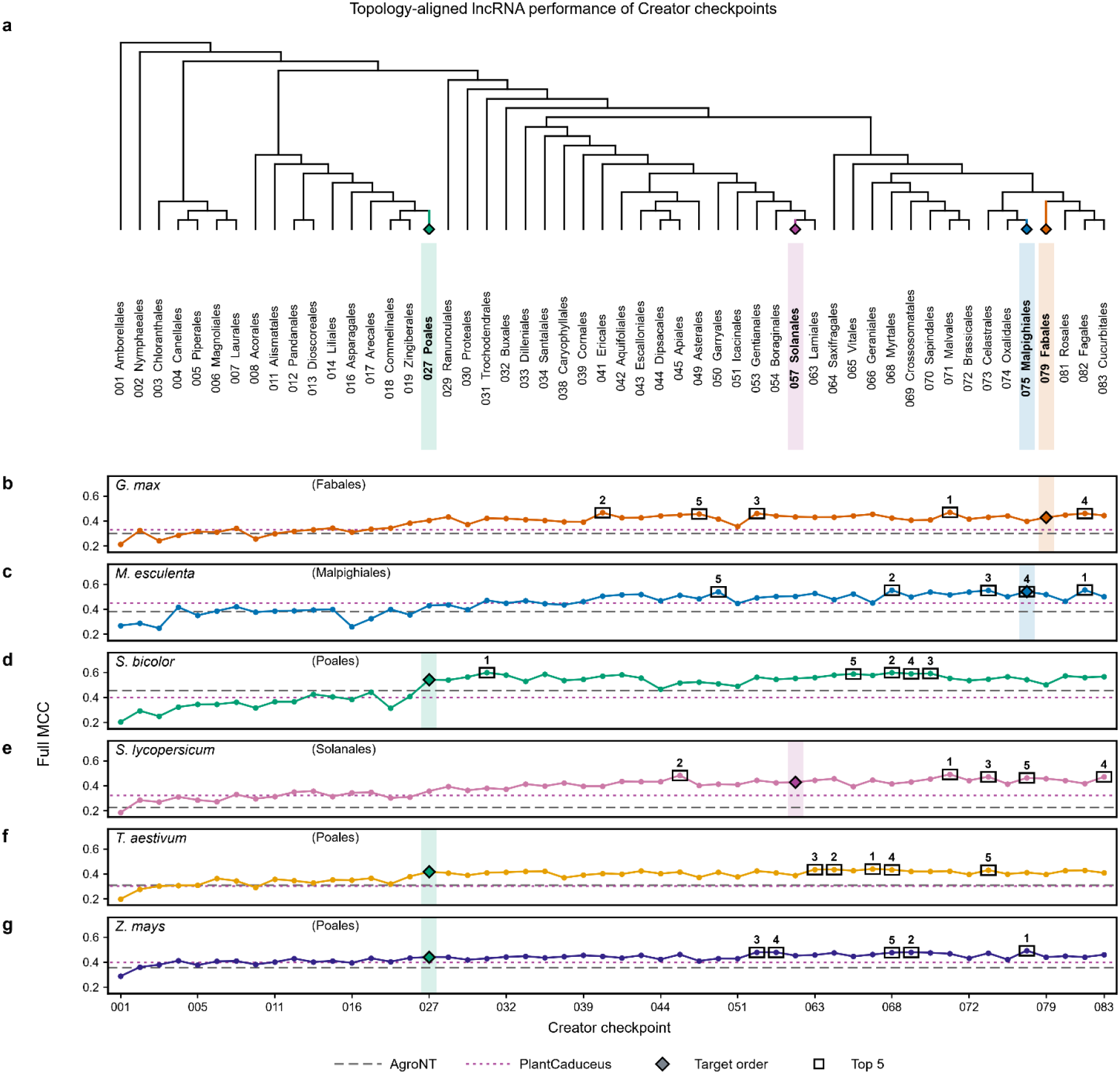
Topology-aligned performance of 52 cumulative Creator checkpoints across six plant lncRNA tasks. a, Rooted phylogenetic topology of 52 plant orders, displayed without branch lengths. The horizontal order follows the first occurrence of each order in the species metadata spreadsheet and is consistent with the cumulative checkpoint sequence. All terminal nodes are aligned. Internal node positions are determined by the number of edges in the retained topology; distances in the tree do not represent evolutionary divergence or time. b–g, Full-data MCC values for 52 checkpoints on lncRNA classification tasks involving soybean, cassava, sorghum, tomato, wheat and maize, respectively. Colored terminal branches, bold black order labels, lightly shaded columns and diamond symbols highlight four target orders and their associated checkpoints. Order labels indicate the taxonomic associations of cumulative checkpoints. Long gray and short purple dashed lines indicate AgroNT and PlantCaduceus baselines, respectively, rerun locally using the same lncRNA evaluation workbook. Black outlined boxes and labels numbered 1–5 identify the five highest-scoring checkpoints for each task, ranked exclusively among the 52 Creator checkpoints using unrounded full-data MCC values.

Stage-dependent performance trajectories also differed among the six tasks (Fig. 5). Under full-data and 10-shot settings, the arithmetic means of the three pairwise Spearman correlation coefficients among the three Poales-related tasks were 0.791 and 0.809, respectively. After adjusting for the rank trend associated with training stage, the corresponding mean partial correlations decreased to 0.554 and 0.587. Checkpoints associated with the taxonomic order of a target species did not consistently yield the highest performance. Across the 18 combinations of six tasks and three supervision settings, only the Poales-associated stage 27 checkpoint ranked first among the 52 Creator checkpoints on the wheat 10-shot task. These findings indicate that checkpoint-associated taxonomic labels do not directly predict optimal performance on tasks involving the corresponding species.

**Fig 5.**
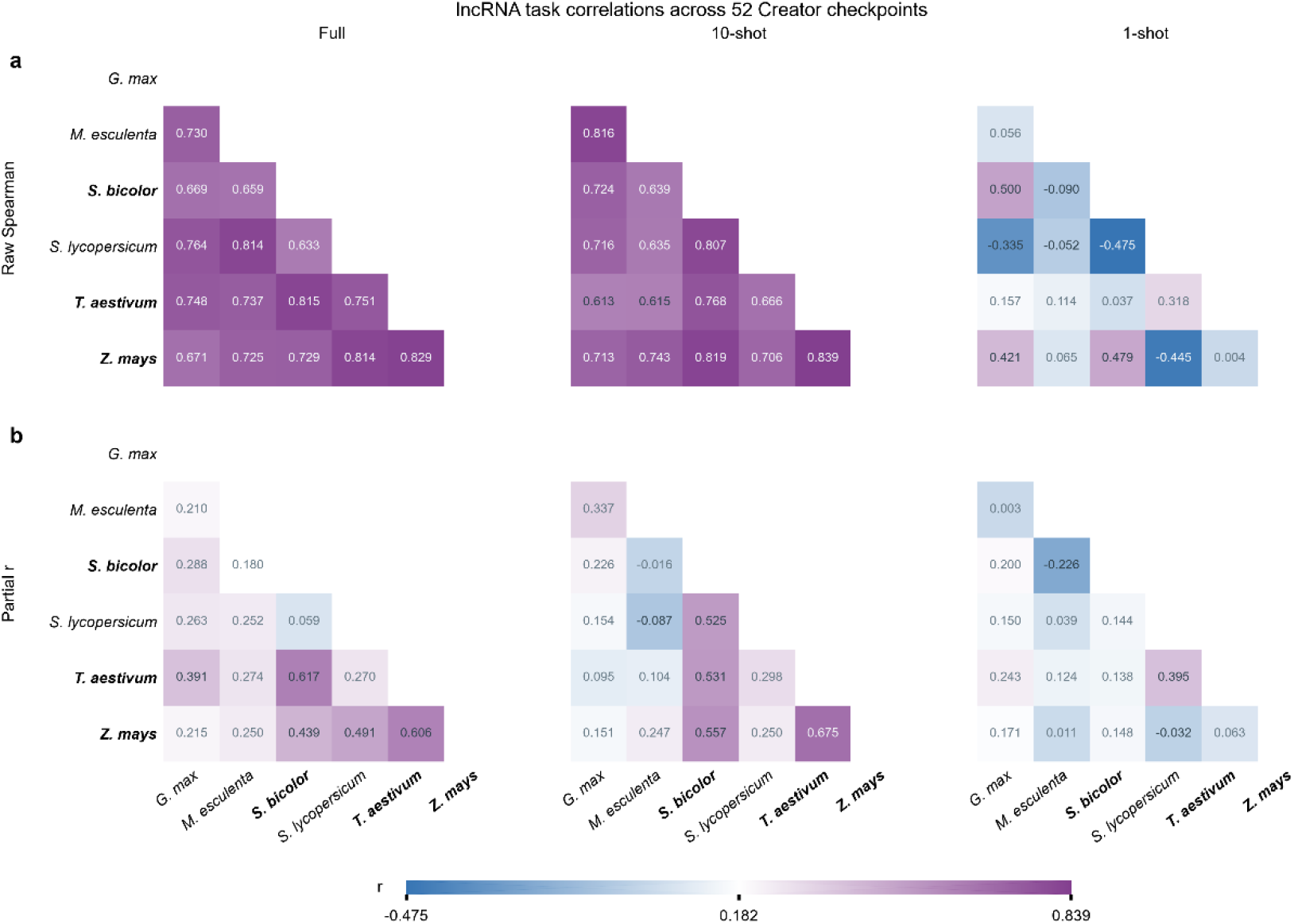
Raw and stage-rank-adjusted correlations among plant lncRNA tasks across 52 cumulative Creator checkpoints. Pairwise correlation matrices of MCC trajectories for six plant lncRNA tasks across 52 cumulative Creator checkpoints. Rows correspond to full-data, 10-shot and 1-shot settings, respectively. a, Raw Spearman correlations. b, Stage-rank-adjusted correlations, calculated as Pearson correlations between residuals obtained by independently regressing the ranked MCC values of each task on an intercept and the ranked stage index using ordinary least squares. Residuals were not ranked again. A total of 90 correlation coefficients are annotated across the six strictly lower-triangular matrices. Names of Poales-related tasks are highlighted in bold black. All six matrices share a common color scale, with white corresponding to the arithmetic midpoint of the displayed coefficient range.

## 3. Discussion

Creator 1.0 was pretrained from random initialization exclusively on plant genomic sequences and achieved the highest mean full-data MCC across the 10 plant-related GENEB tasks, while ranking among the top-performing models across all 100 tasks. The stage 81 checkpoint also achieved the highest scores on selected non-plant 4mC classification tasks. However, performance varied across low-data settings, target species and score aggregation methods, indicating that overall benchmark scores alone are insufficient to characterize model applicability across different task categories.

Evaluation of 52 cumulative checkpoints further revealed task- and supervision-dependent differences in performance on plant lncRNA classification tasks. These findings suggest that phylogeny-guided stagewise pretraining produces checkpoints with distinct task-dependent representation capabilities, providing an additional basis for model selection in plant genomic applications. Although the current results do not establish a consistent advantage of lineage-associated checkpoints for their corresponding species, further refinement of training stages, task-specific adaptation and independent validation may enable the development of specialized models for different plant orders, families, genera and individual species. Such an approach could provide more targeted model resources for plant genome functional analysis and related applications.

## 4. Materials and Methods

### 4.1. Pretraining corpus and model configuration

Genomic sequences from 1,000 plant species were compiled from public databases ^13^ and organized into 83 consecutive training stages according to a fixed phylogenetic tree. At each stage, training continued from the preceding model parameters, with checkpoints retained alongside lineage and training-batch metadata. The reported 1.11 Tbp represents the aggregate genome assembly size calculated from source records. Creator 1.0 was trained from random initialization exclusively on plant genomic sequences using self-supervised learning. The model was based on the Gemma 3 architecture ^14^, comprising approximately 2 billion parameters, 30 Transformer layers, a hidden dimension of 2,048, eight query attention heads, two key–value heads and an intermediate dimension of 9,216. DNA sequences were encoded using non-overlapping 3-mers, with a maximum input length of 32,768 tokens. The saved training configuration specified BF16 precision, the AdamW optimizer, a learning rate of 2 × 10^−4^, a cosine learning-rate schedule and a warm-up ratio of 3%.

### 4.2. GENEB benchmark evaluation

Six checkpoints (stages 27, 49, 63, 72, 81 and 83) were evaluated across 100 GENEB classification tasks. The Creator backbone was frozen, sequence representations were extracted, and only logistic regression probes were fitted. We used the official data splits and full-data, 10-shot and 1-shot evaluation settings, where *shot* denotes the number of training examples per class. Evaluations were conducted using five fixed random seeds (13, 17, 42, 123 and 997), with a batch size of 8 for representation extraction. For each task, MCC values were first averaged across the five seeds, followed by equal-weight averaging across tasks. For category-balanced aggregation, MCC values were first averaged within each functional category and then equally averaged across the 13 categories. All model rankings were determined using unrounded scores. Results for the 40 publicly available reference models were obtained from the official GENEB benchmark and were not independently rerun in this study ^9^. Details of the evaluation code, data versions and task sources are provided in the Supplementary Information.

### 4.3. Plant lncRNA tasks and stage-specific checkpoint evaluation

We evaluated 52 Creator checkpoints on six binary lncRNA classification tasks involving soybean, cassava, sorghum, tomato, wheat and maize ^9^, using the same frozen-representation and linear-probing framework. AgroNT (agro-nucleotide-transformer-1b) ^6^ and PlantCaduceus (PlantCaduceus_l32) ^8^ were independently rerun locally using identical data splits, random seeds and supervision settings. MCC was used as the primary evaluation metric. We calculated both the equally weighted mean MCC across the six tasks and the equally weighted mean across four target plant orders. For the latter, task-level MCC values were first averaged within each order, followed by equal-weight averaging across the four orders. Rankings of target-order-associated checkpoints were calculated exclusively among the 52 Creator checkpoints for each task. Details of the execution environments and model versions used across evaluation runs are provided in the Supplementary Information.

Spearman rank correlations were used to assess similarities in stage-dependent performance trajectories between tasks, with average ranks assigned to tied values. To calculate stage-rank-adjusted partial correlations, the ranked MCC values for each task were regressed on an intercept and the ranked stage index using ordinary least squares. Pearson correlations were then calculated between the resulting task-specific residual vectors, without further ranking of the residuals. For each supervision setting, only the 15 unique task pairs in the strictly lower-triangular correlation matrix were displayed, yielding 90 coefficients across three supervision settings and two correlation methods.

## Code and Data Availability

The source code, genomic data provenance records and model files are planned for public release following completion of further validation. Corresponding repository and access links will be provided upon release.

## References

1. Zhou, Z. et al. in International Conference on Learning Representations, Vol. 2024 41642–41665 (2024).

2. Nguyen, E. et al. Hyenadna: Long-range genomic sequence modeling at single nucleotide resolution. Advances in neural information processing systems 36, 43177–43201 (2023).

3. Schiff, Y. et al. Caduceus: Bi-directional equivariant long-range dna sequence modeling. Proceedings of machine learning research 235, 43632 (2024).

4. Sanabria, M., Hirsch, J., Joubert, P.M. & Poetsch, A.R. DNA language model GROVER learns sequence context in the human genome. Nature Machine Intelligence 6, 911–923 (2024).

5. Dalla-Torre, H. et al. Nucleotide transformer: building and evaluating robust foundation models for human genomics. Nature methods 22, 287–297 (2025).

6. Mendoza-Revilla, J. et al. A foundational large language model for edible plant genomes. Communications Biology 7, 835 (2024).

7. Benegas, G., Batra, S.S. & Song, Y.S. DNA language models are powerful predictors of genome-wide variant effects. Proceedings of the National Academy of Sciences 120, e2311219120 (2023).

8. Zhai, J. et al. Cross-species modeling of plant genomes at single-nucleotide resolution using a pretrained DNA language model. Proceedings of the National Academy of Sciences 122, e2421738122 (2025).

9. Ledneva, D., Nuridinov, M. & Kuznetsov, D. GENEB: Why Genomic Models Are Hard to Compare. arXiv preprint 2606.04525 (2026).

10. Marin, F. et al. in International Conference on Learning Representations, Vol. 2024 15246–15281 (2024).

11. Patel, A. et al. Dart-eval: A comprehensive dna language model evaluation benchmark on regulatory dna. Advances in Neural Information Processing Systems 37, 62024–62061 (2024).

12. Feng, H. et al. Benchmarking DNA foundation models for genomic and genetic tasks. Nature communications 16, 10780 (2025).

13. Database resources of the national genomics data center, China national center for bioinformation in 2025. Nucleic acids research 53, D30–D44 (2025).

14. Team, G. et al. Gemma 3 technical report. arXiv preprint 2503.19786 (2025).

